# Interpretable XGBoost-SHAP model predicts the nanoparticles delivery and reveals its interaction with tumor genomic profiles

**DOI:** 10.1101/2022.06.06.494964

**Authors:** Yuxia Tang, Xingqun Ma, Shouju Wang

**Affiliations:** Laboratory of Molecular Imaging, Department of Radiology, The First Affiliated Hospital of Nanjing Medical University, Nanjing, Jiangsu, China; Department of Oncology, Nanjing Baiyi Hospital, Jinling Clinical College of Nanjing Medical University, Nanjing, Jiangsu, China

**Author notes:** Corresponding author: Shouju Wang.

## Abstract

Understanding the complex interaction between nanoparticles (NPs) and tumors *in vivo* and how it dominates the delivery efficacy of NPs is critical for the translation of nanomedicine. Herein, we proposed an interpretable XGBoost-SHAP model by integrating the information of NPs physicochemical properties and tumor genomic profile to predict the delivery efficacy. The correlation coefficients were > 0.99 for all training sets, and 0.830, 0.839, and 0.741 for the prediction of maximum delivery efficacy (DEmax), delivery efficacy at 24 h (DE24), and delivery efficacy at 168 (DE168) for test sets. The analysis of the feature importance revealed that the tumor genomic mutations and their interaction with NPs properties played an important role in the delivery of NPs. The functional profile of the NP-delivery-related genes was further explored through gene ontology enrichment analysis. Our work provides a method to accurately predict the delivery efficacy of NPs to heterogeneous tumors and highlights the power of simultaneously using omics data and interpretable machine learning algorithms for discovering the interaction between NPs and tumors, which is important for the development of precision nanomedicine.

## Introduction

Nanoparticles (NPs) provide a promising platform for the diagnosis and treatment of tumors as they can be engineered to have multiple functions and excellent physicochemical properties. ^[1]^ There have been several NPs successfully translated to clinics for tumor imaging and treatment, such as ^64^Cu-MM-302 and Doxil. ^[2,3]^ These NPs were designed to selectively deliver to tumors through the enhanced penetration of retention (EPR) effect and thus reduce the background signal or side effects. However, recent meta-analyses reveal the delivery efficacy is low for most published NPs and tumor models, ^[4,5]^ which underpins the importance of understanding the complex interaction between NPs and tumors *in vivo.*

Numerous efforts have been devoted to exploring the relationship between the physicochemical properties and pharmacokinetic profile of NPs. Early studies have shown that the size, stiffness, shape, and surface modification would dictate the delivery efficacy of NPs. ^[6–9]^ Very recently, several reports applied machine learning (ML) and literature data mining techniques to predict the biodistribution of NPs *in vivo* from the combination of their properties. ^[10,11]^ These ML models reflect the complex relationship between the delivery efficacy and the properties of NPs, which is not possible for a single study based on a small number of NPs. ^[12]^ Nonetheless, it remains challenging to use these ML models to explain the difference in biodistribution among different tumor models and how much an NP property influences the final delivery efficacy in a certain tumor.

Although the influence of NPs properties has been extensively studied, the impact of biological heterogeneity of tumors on the delivery efficacy has been overlooked for a long time. Most NPs made in labs were only tested on a few tumor models with the same cell line and similar size. Clinical reports have shown that the tumor characterization, such as size, type, angiogenesis, and perfusion, would greatly affect the EPR magnitude and thus the delivery efficacy of NPs. ^[13–15]^ Experience with small molecular drug discovery also suggested that the genomic heterogeneity of tumors helps identify drug targets and patient selection. For example, the omics databases of NCI-60 and Cancer Cell Line Encyclopedia (CCLE) have been widely used to screen small molecular drugs and validate tumor targets. ^[16,17]^ Therefore, we hypothesize integrating pan-cancer genomics profile and NPs properties as features into the ML models would help to elucidate the complex interaction between NPs and tumors *in vivo* and predict the delivery efficacy more accurately.

The interpretability of ML models is critical for understanding the mechanism behind the complex interaction between the physicochemical properties of NPs and the biological heterogeneity of tumors. Tree-based ensemble ML models, e.g., eXtreme Gradient Boosting (XGBoost) and random forest (RF), are usually comparable in performance and more interoperable than neural networks. The interpretability of these ML models is further improved by the techniques like Shapley addictive explanations (SHAP). ^[18,19]^ SHAP explores and illustrates the importance of and correlations between input features based on the game-theoretic approach, which has been applied to explore the gene-gene and gene-environment interactions very recently. ^[20]^

To capture the NP-tumor interaction complexities and enable the accurate prediction of delivery efficacy of NPs in different tumors models, we developed ML models by integrating NPs properties of 162 samples from literature data-mining and genomic profiles of 23 cell lines from the CCLE database. We observed that the genomic mutations of tumors have a dominating influence on the delivery efficacy, which was comparable to the influence of tumor weight and NPs properties. Moreover, through the analysis of SHAP values, we identified the interaction between genomic mutations and NP properties, and how they determined the prediction of delivery efficacy together. We further analyzed the molecular function, biological process, protein type, and pathway of the related genes through gene ontology (GO) enrichment analysis. The models proposed here revealed the value of omics data for explaining the NP-tumor interaction complexities, filling the long-missing piece of the study of precision nanomedicine.

## Materials and Methods

### Datasets and data preprocessing

The delivery efficacy dataset was obtained from the previously published meta-analysis, ^[10]^ containing the delivery efficacy at 24 h (DE24), 168 h (DE168), and the maximum delivery efficacy (DEmax) after intravenous injection of NPs. The original dataset contains 330 samples. The samples tested on non-human tumor cell lines (146 samples) and cell lines not included in the CCLE database (22 samples) were excluded. Finally, 162 samples covering 23 tumor cell lines were included in the following analysis.

To describe the variables of samples, six NPs properties [*i.e*., nanomaterials (polymersome, liposomes, dendrimers, hydrogels, other organic materials, gold, silica, iron oxide, and other inorganic materials), types (organic and inorganic), shape (spherical, rod, and other shapes), targeting strategy (active and passive), size and zeta potential] and three tumor characteristics [*i.e*., tumor weight, cell lines and models (heterotopic and orthotopic)] were extracted from the publications by hand. Some mislabeled cell lines (KB and Bcap37) were corrected to HeLa. The details are shown in supplementary Table 1. The distribution of the dataset was visualized by tabplot package in R (version 4.2.0). ^[21]^

To characterize the biological heterogeneity of tumor cell lines, the genomic mutation data of the 23 tumor cell lines were extracted from the CCLE database. ^[17]^ The gene status was encoded as a binary variable (mutated: 1, widetype: 0). The conservative mutations were excluded because they result in amino acids with similar properties. The mutations that were not hotspots in The Cancer Genome Atlas (TCGA) database were also excluded. Only the genes mutated at least once in the 23 cell lines were kept for the following feature selection (m = 277). To further exclude the redundant and irrelevant mutations, Least Absolute Shrinkage and Selection Operator (LASSO) regression was performed using the 277 genes as input and the DE24, DEmax, and DE168 as output, respectively. Finally, 63 genes were selected for the prediction of DE24, 66 genes for DEmax, and 60 genes for DE168. The selected genes and corresponding mutant status were joined with the delivery efficacy dataset according to the index of tumor cell lines.

To increase the speed of model training, the numerical variables (size and zeta potential) were standardized by the following formula.

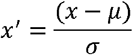

where *x’* is the standardized feature value, *x* is the feature value, *μ* is the expectation of the feature, and *σ* is the standard deviation of the feature. Missing values were filled by the mean value of all other samples. One-hot encoding was performed on nominal variables (NPs materials and shape) to avoid assuming orders of these variables.

### Machine learning regression and validation

Five ML algorithms were applied to establish models for the prediction of delivery efficacy, including XGBoost, random forest (RF), light gradient boosting machine (LGBM), support vector machines (SVM) with radial basis function (RBF) kernel or linear kernel. The number of estimators was set as 500 for XGBoost, RF, and LGBM. The algorithms were implemented in Python 3.8 with packages including xgboost (version 1.6.1), scikit-learn (version 1.1.1), and lightgbm (version 3.3.2). The models were constructed with and without genomic mutation information. Random search method was applied to tune the hyperparameters if needed.

The original dataset was split into a training set (80%) and a test set (20%). The performance of models was evaluated by the coefficient of determination (R^2^) and mean square error (MSE) between the predictions and observations of test sets. To prevent overfitting, 10-fold cross-validation with shuffle split was applied to each ML model. The calculation of R^2^ and MSE was defined as below:

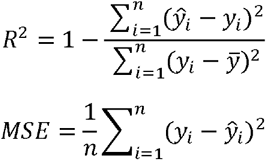

where *y_i_* and *y*□*_i_* are the observed and predicted delivery efficacy, *y*□ is the mean of observed delivery efficacy, and *n* is the total number of samples. A higher value of R^2^ and a lower value of MSE suggest better fitness of the ML model.

### Feature importance analysis of XGBoost models

To interpret the feature importance of XGBoost models, the SHAP approach was introduced to the models. A unique SHAP value was assigned to each feature of each sample to represent the deviation from the average prediction brought by the value of a feature in one sample. A positive SHAP value indicates the value of a feature increases the probability of higher delivery efficacy and *vice versa.* SHAP is the only explainer that stratifies local accuracy, missingness, and consistency for ML model interpretation. The SHAP value was calculated by the SHAP package (version 0.4.0) using the formula below:

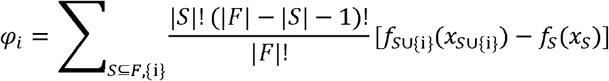

where *f* represents the prediction model, *F* represents the set of all features and *S* represents the subset of all features excluding the *i*^th^ feature.

The average absolute SHAP values (mean of |SHAP|) of each feature were used to measure the feature importance. The sum of the average |SHAP| of all genes was used to represent the importance of genomic mutations in the models. The dependence plots of SHAP values were used to illustrate the relationship between NPs properties and their influence on the prediction of delivery efficacy in tumor cell lines with or without certain gene mutations.

### GO enrichment analysis

To further identify the functional profile of the selected genes in the ML models, GO enrichment analysis was performed on the PANTHER online tool ((http://pantherdb.org/, version 17). ^[22]^ The commonly shared genes in the models of DE24, DEmax, and DE168 were input into the PANTHER tool and the molecular function, biological process, protein class, and pathway analyses were performed using the default settings.

## Results

The pipeline of literature data mining, genomic profile integration, model development, and feature analysis was depicted in Figure 1 with specifics as provided in Materials and Methods. Based on the meta-analysis conducted by Lin’s group, a total of 162 samples with six NPs properties (nanomaterials, types, shape, targeting strategy, size, and zeta potential) and three tumor characteristics (tumor weight, cell lines, and models) were extracted (Figure 2A). Twenty-three cell lines covering 12 lineages (breast, cervix, central nervous system, pancreas, colorectal, gastric, liver, lung, lymphocyte, ovary, prostate, skin) were included. Further details of the information extracted from the literature were listed in supplementary Table 2.

**Figure 1.**
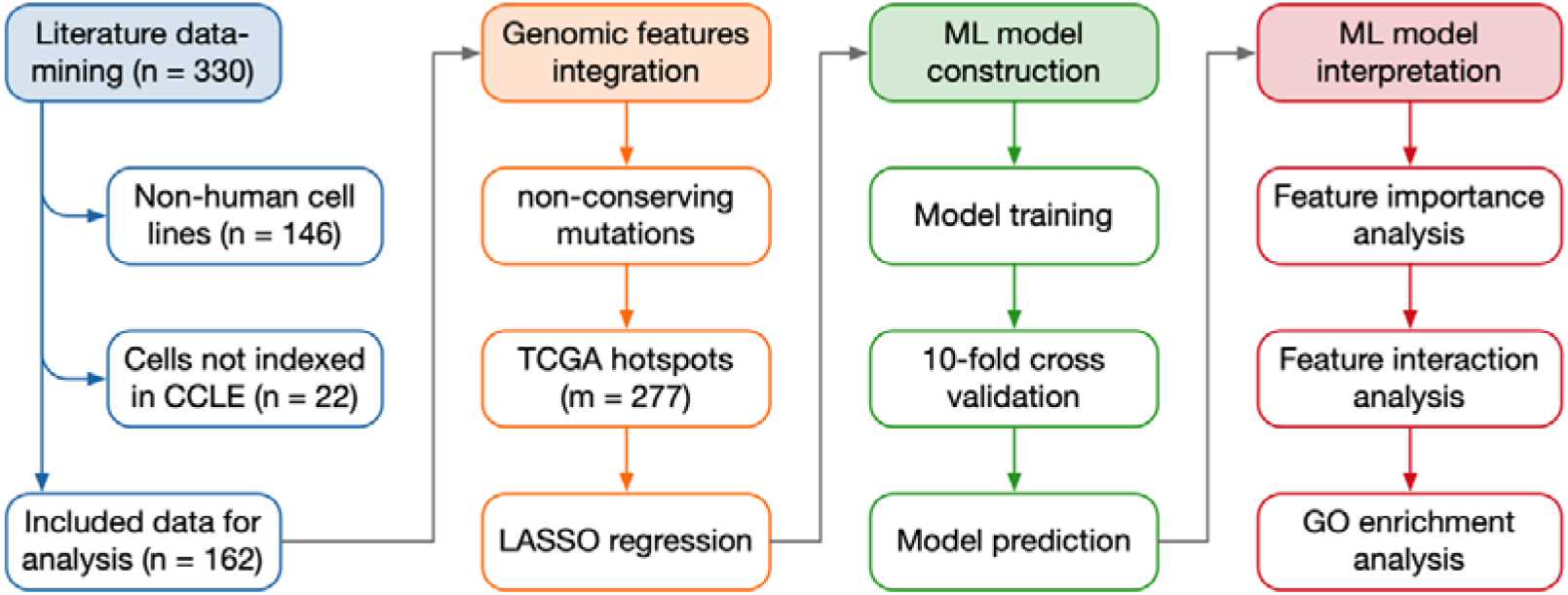
Pipeline for the literature data-mining, genomic features integration, ML model construction and interpretation. CCLE: Cancer Cell Line Encyclopedia; TCGA: The Cancer Genome Atlas; ML: Machine Learning; GO: Gene Ontology.

**Figure 2.**
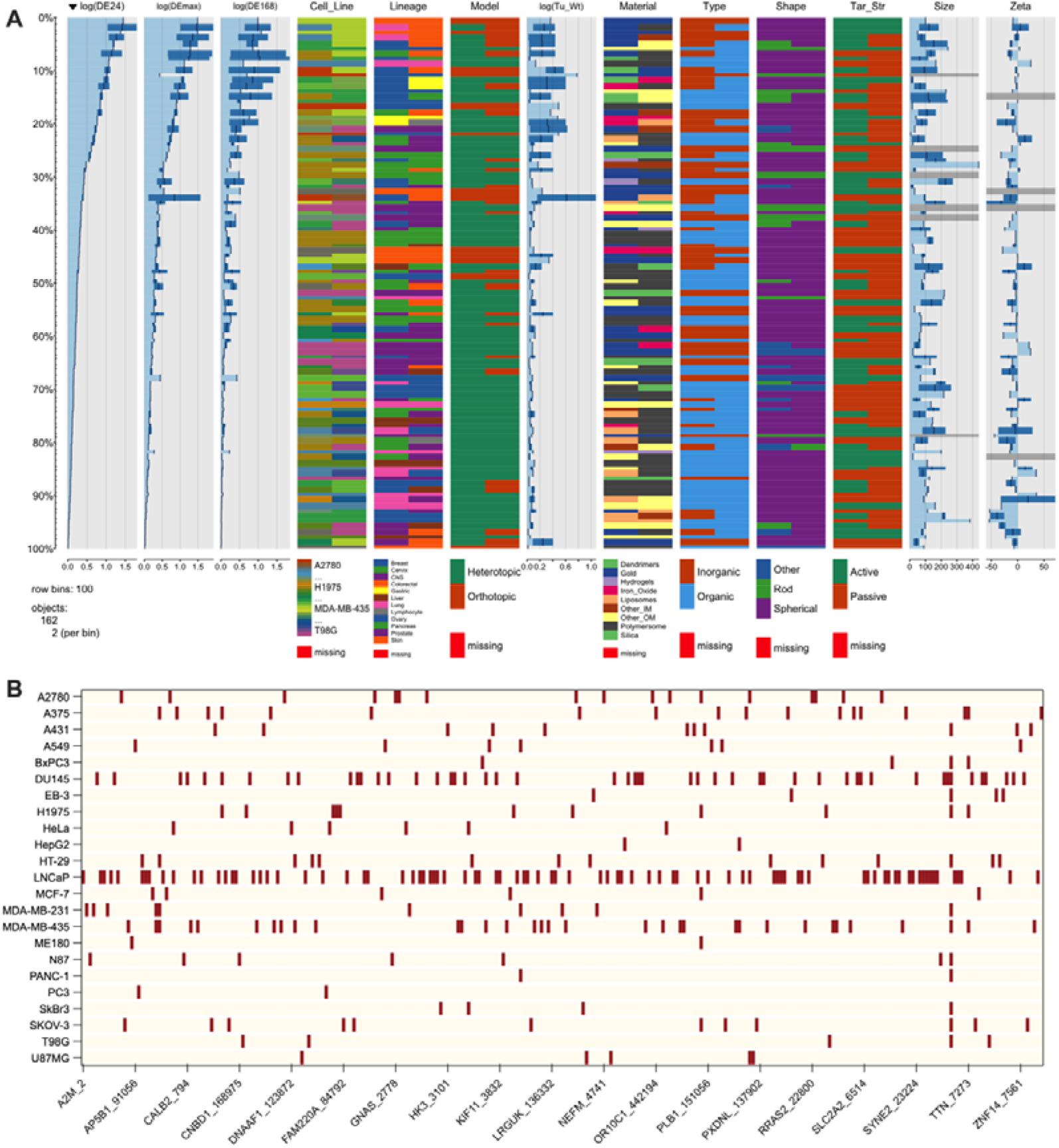
Visualization of the delivery efficacy dataset consisting of 162 samples and the genomic profile of the 23 tumor cell lines. (A) The delivery efficacy dataset visualized by ‘tabplot’ package in R. The dataset is split into 100 bins and sorted according to the values of DE24. Logarithmic transformation is performed for DE24, DEmax, DE168, and tumor weight. Tu_Wt: Tumor weight; Tar_Str: target strategy. (B) Heatmap of genomic profile of each tumor cell line. The brown bars indicate mutant genes. The genes are represented as ‘HUGO name_ NCBI Entrez ID’.

The genomic mutation information was extracted from the CCLE database. To increase the predictive power of the ML models, the mutation information underwent rigid feature selection before being integrated with the delivery efficacy dataset. The mutations that were conservative or not hotspots in the TCGA database were excluded. The remaining 277 genes were depicted in Figure 2B and underwent LASSO regression to remove redundant and irrelevant genes for delivery efficacy prediction. Overall, 63, 66, and 60 genes were selected to construct the ML models for the prediction of DE24, DE168, and DEmax. The detail of selected genes was listed in supplementary Table 3. Figure 2 shows no observable trend in delivery efficacy as a function of NPs properties or tumoral genomic mutations, indicating the necessity of further analysis with flexible ML models.

Ten ML models were constructed to predict the delivery efficacy of NPs with or without tumor genomic profiles. The R^2^ values of the ten models were summarized in Table S1-3. It is shown that the XGBoost models trained with tumor genomic profiles have the best performance in 10-fold cross-validation and testing set prediction. The R^2^ values of XGBoost models trained with genomic mutation information for DE24, DEmax, and DE168 prediction were 0.997, 0.997, and 0.994 for the training sets, and 0.839, 0.830, and 0.741 for the testing sets (Figure 3). It is noted that the models trained with genomic mutation information have better performance than those trained without that information, indicating that the genomic mutation information is critical for the accuracy of prediction *via* ML models.

**Figure 3.**
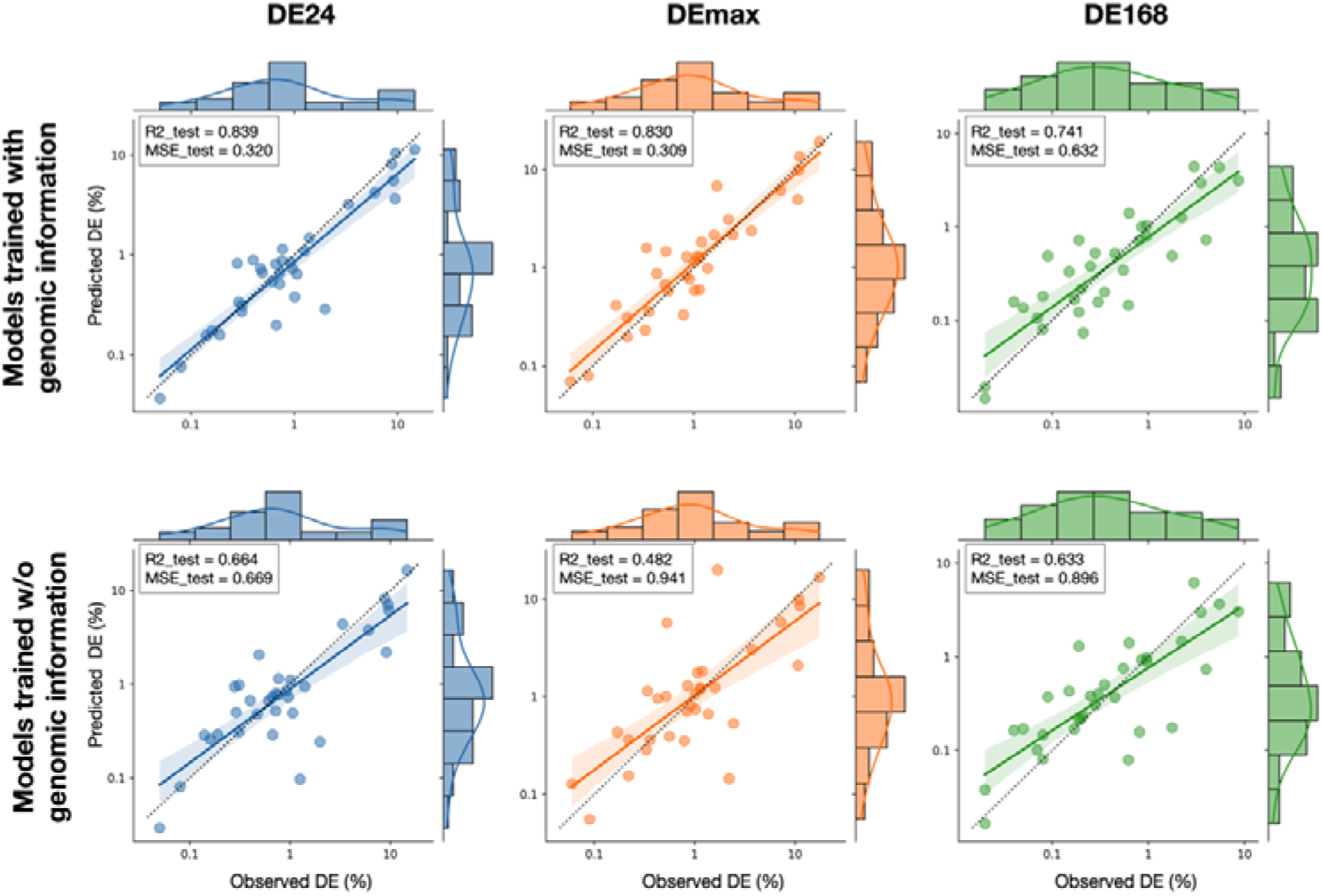
Prediction performance of the XGBoost models for test sets. The upper row summarizes the performance of models trained with the genomic information for the prediction of DE24, DEmax, and DE168. The lower row summarizes the performance of models trained without the genomic information. The R2 and MSE values are depicted in the legends. DE: delivery efficacy.

To interpret the influence of features (NPs properties and tumor characteristics) on the delivery efficacy, the SHAP algorithm was applied to measure the feature importance in the XGBoost models. Figure 4A-C depicted the summary and beeswarm plots of the top 17 features for the prediction of DE24, DEmax, and DE168. The summary plots show that the tumor weight, as well as the size and zeta potential of NPs, are the most important three features that dominate delivery efficacy. The mutation of certain genes, such as DNAAF1 and ATP10A, also influenced remarkably on the output. The beeswarm plots show that heavier tumor weight and mutation of DNAAF1 and ATP10A result in higher SHAP values, which indicate an increased probability of efficient delivery. In contrast, lighter tumor weight and the mutation of NRAS result in lower SHAP values, which indicate a decreased probability of efficient delivery. The impact of NPs size and zeta potential on prediction seems more complex. This result is in line with previous clinical observations and the function of related genes. It is reported that patients with larger tumors often exhibited a greater magnitude of EPR effect. ^[13]^ The DNAAF1 involves in the regulation of microtubules and is related to the uptake of lipoprotein. ^[23]^ The ATP10A belongs to the family of P-type cation transport ATPases and is related to the ATPase-coupled cation transmembrane transporter activity. ^[24]^

**Figure 4.**
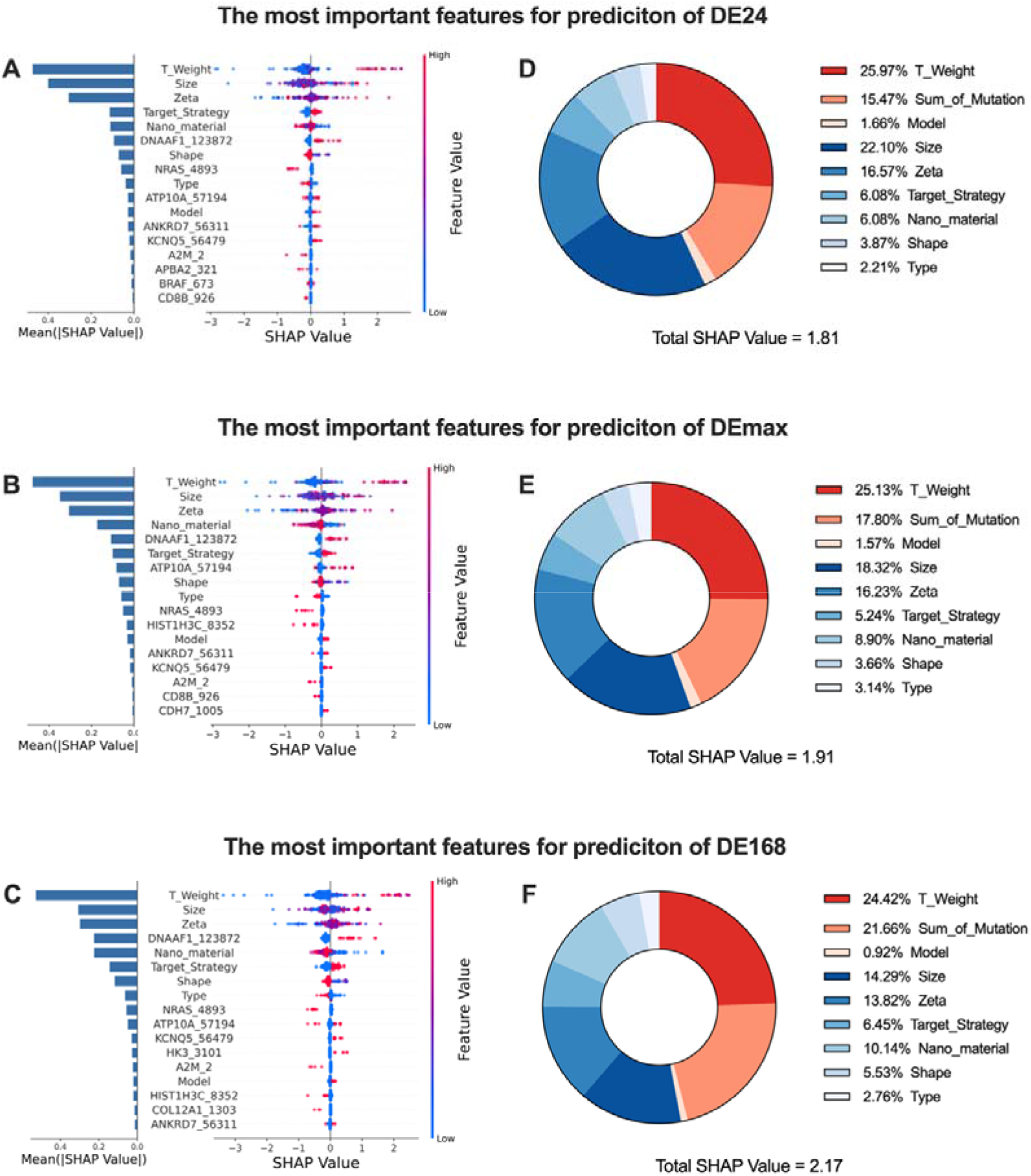
Feature importance analysis by SHAP. (A-C) The summary plots (left column) and beeswarm plots (right column) of the most important 17 features for the prediction of DE24, DEmax, and DE168. Each row represents a feature. The genes are represented as ‘HUGO name_ NCBI Entrez ID’. (D_F) The donut plots showing the proportion of influence from various features. T_weight: Tumor weight. The tumor characteristics are represented in warm hue. The NPs properties are represented in cold hue.

To compare the influence of tumor heterogeneity and NPs properties on delivery efficacy, the proportion of features’ SHAP values was illustrated in Figure 4D-E. It is shown that the biological heterogeneity of tumors (including tumor weight, genomic mutations, and model type) contributed almost half of the total SHAP value of features (43.1% for DE24 prediction, 44.55% for DEmax prediction, and 47.00% for DE168 prediction). Noted the genomic mutations contributed roughly 20% of the total SHAP value (15.47% for DE24 prediction, 17.80% for DEmax prediction, and 21.66% for DE168 prediction), which is comparable with the influence of the size or zeta potential of NPs, suggesting the importance of genomic mutations to the delivery efficacy of NPs *in vivo.*

To further explore how genomic mutations affect the influence of NPs properties on delivery efficacy, SHAP dependence plots were depicted in Fig 5. The x-axis represents the values of the NPs properties (size or zeta potential), and the y-axis represents the corresponding SHAP values. Orange dots indicate samples with mutated genes, while blue dots indicate samples with wide phenotypes. It is shown that the SHAP values increase with larger sizes of NPs for PTEN-mutated tumors, which suggests that increasing NP size increases the probability of efficient delivery to these tumors. In contrast, for PTEN-widetype tumors, the SHAP values tend to decrease with larger NPs sizes, which indicates the same NPs properties may have opposite influence on delivery efficacy depending on the genomic profile of tumors. Similarly, the mutation of ATP10A exhibited interaction with the zeta potential of NPs for the prediction of DE24, DEmax, and DE168. The delivery efficacy to ATP10A mutated tumors showed higher sensitivity to the change of zeta potential, which is probably because ATP10A controls the activity of transmembrane cation transporters. ^[24]^ These results indicate the optimization of NPs should consider the genomic profile of tumors and its interaction with NPs properties, which has been suggested in the concept of precision medicine but neglected long in the development of nanomedicine.

**Figure 5.**
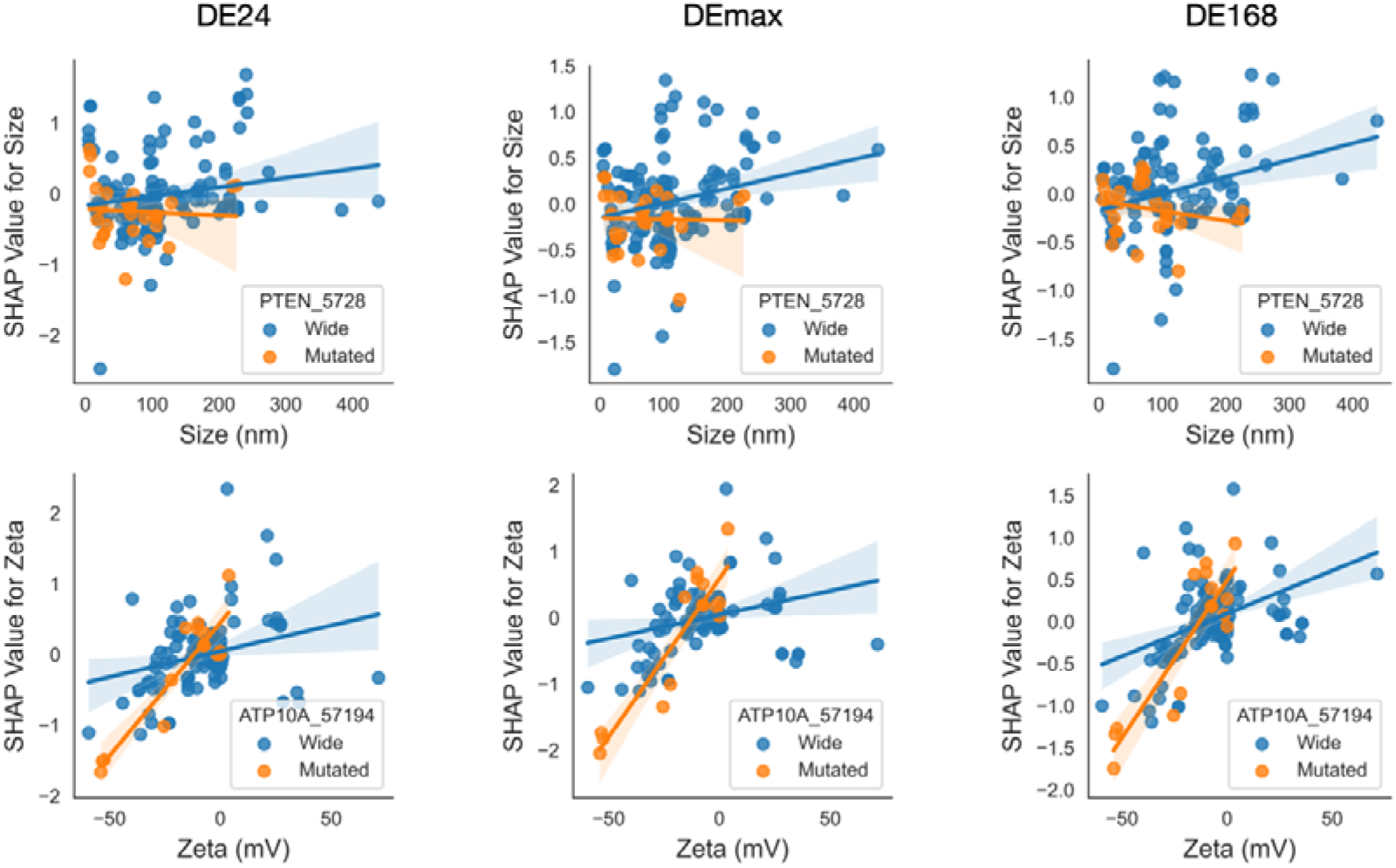
The dependence plots of SHAP values calculated from XGBoost models of DE24, DEmax, and DE168. The x axis represents the value of NPs properties (size or zeta potential). The y axis represents the corresponding SHAP values of the NPs property. Orange dots represent samples tested in tumors with mutant PTEN or ATP10A. Blue dots represent samples tested in widetype tumors.

To gain insight into the function of selected genes as a whole, the commonly shared genes for prediction were identified and underwent GO enrichment analysis using PANTHER online tools. ^[22]^ Figure 6 shows that 56% (45/80) of the genes were shared by the three XGBoost models for delivery efficacy prediction. The molecular functions of these genes involved binding, catalytic activity, transporter activity, and molecular transducer activity. The biological processes of these genes were related to the metabolism, signaling, and response to stimulus. Most of these genes coded proteins related to cell uptake, such as transporters, scaffold/adaptor proteins, transmembrane signal receptors, and protein-binding activity modulators. The pathway analysis showed these genes participate in angiogenesis, inflammation, EGF receptor- and integrin-signaling, which is highly related to the EPR effect of tumors. ^[25]^

**Figure 6.**
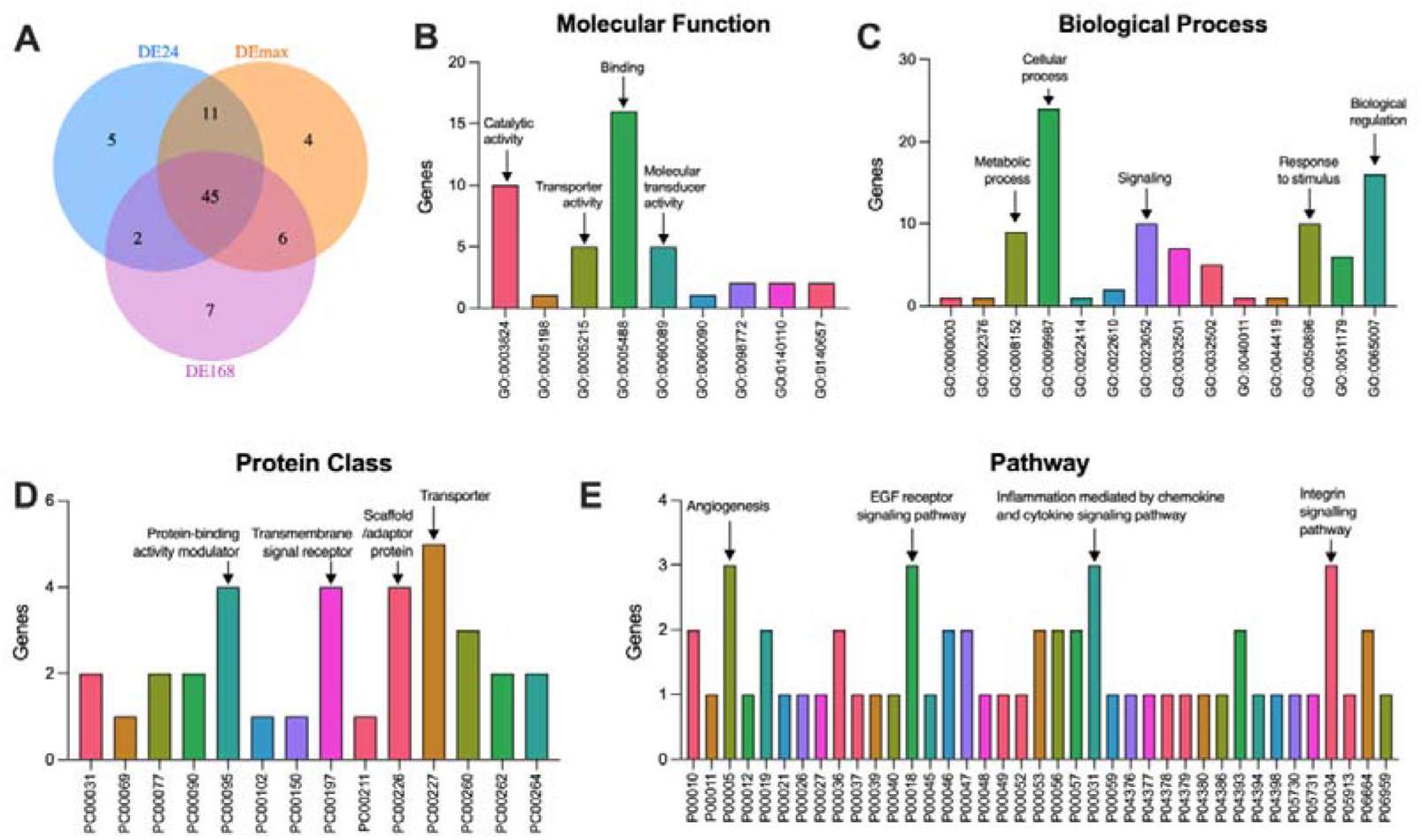
The GO enrichment analysis of delivery-efficacy-related genes. (A) The Venn diagram showing the shared genes in the XGBoost models for the prediction of DE24, DEmax, and DE168. (B-E) the molecular function, biological process, protein class, and pathway of the shared genes analyzed by PANTHER online tools.

## Discussion

In this work, we represent the first effort of integrating NPs properties and tumor genomic profile to predict the delivery efficacy of NPs *in vivo* by ML models. The best models were based on XGBoost with R^2^ values higher than 0.99 for the training sets and reached 0.839, 0.830, and 0.741 for the testing set in the prediction of DE24, DEmax, and DE168. Taking advantage of the SHAP algorithm and GO enrichment analysis, the importance of and the complex interactions between NPs properties and tumor genomic profile were interpreted, which has not been explored in previous studies.

Our analysis highlights the strong influence of tumor heterogeneity on the delivery efficacy of NPs. Previous studies have suggested that the tumor size is related to the EPR magnitude of tumors, which is revealed in our models as the positive relationship between delivery efficacy and tumor weight. ^[13]^ The tumor type was reported as another important determinant in predicting delivery efficacy, ^[10,13]^ which is also confirmed by the importance of genomic profile in our models. Because there is strong multicollinearity between tumor types/lineages and the genomic profiles, we excluded the tumor types/lineages from the model construction to increase the reliability of the statistical inferences. Another advantage of using genomic profiles instead of tumor types/lineages is that some mutations are shared across cell lines, which is beneficial to explore the common mechanism that dominates the delivery efficacy through unregulated signaling pathways.

One strength of our approach is the application of interpretable ML models, such as XGBoost and SHAP, allowing us to explain how a single feature affected the prediction of delivery efficacy. Since the SHAP values were calculated for each feature of each sample, we could estimate the interaction between NPs properties and tumor heterogeneity, *i.e.,* the exact influence of changing one physicochemical property of an NP formula on the delivery efficacy to a certain type of tumor. This may serve as a powerful tool for personalized nanomedicine because studies using different models often show opposite trends concerning the optimal NP shape and size for tumor delivery. ^[1]^ Recent studies also showed that the genetic mutation would affect the tumor-targeting ability of certain NPs. For example, albumin-bound NPs exhibited significantly higher delivery efficacy than dextran NPs to KRAS-mutant lung tumors but not to widetype tumors, ^[26]^ which underpins the importance of studying the interactions between NPs properties and tumor heterogeneity.

We noticed the pioneering work to explore the influence of biological heterogeneity on NP-cell interaction through the lens of multi-omics from other groups. ^[27]^ The high-throughput screening used in that study is indeed a powerful tool to identify potential biomarkers for the development of precision nanomedicine. However, the delivery of NPs *in vivo* is dependent on many other factors beyond the NP-cell interactions *in vitro.* For example, the angiogenesis and inflammation pathways were unlikely to be identified as the driving force of efficient delivery at the level of NP-cell interaction analysis.

There are several limitations of our study despite the accuracy and interoperability of the models. First, our model only included the genomic mutation data. It is because other omics data, such as epigenomics, transcriptomics, proteomics, metabolomics, and radiomics, are extremely variable among tumors *in vivo,* and could not be confidently extrapolated from the current annotated datasets. Second, only 23 tumor cell lines were analyzed, and the proportion of cell lines was imbalanced, which might hamper the accuracy of the final models. This problem will be alleviated in the future with the increasing number of tumor cell lines in published papers and the development of data handling techniques like label distribution smoothing. Third, the cell line-derived xenograft models cannot fully capture the heterogeneity of human tumors. We believe valuable insight may be gleaned by analyzing the NPs delivery efficacy with multi-omics data of patients treated by nuclide-tagged NPs following a similar pipeline.

## Conclusions

Overall, this work suggests that powerful and interpretable ML models can be established by integrating information derived from literature data-mining and genomics datasets to predict the delivery of NPs *in vivo.* Similar approaches should be applicable to predict the therapeutic, immune-regulating, and gene-editing efficacy of NPs to tumors. Our work highlights the importance of tumor genomic profile for the behavior of NPs, which is insightful not only for the design of NPs but also for the patient selection in the ear of precision nanomedicine.

---

**Table S1.** R^2^ values for the prediction of DE24 using different ML algorithms.

|  | With mutation information |  |  | Without mutation information |  |  |
| --- | --- | --- | --- | --- | --- | --- |
|  | 10-fold CV | Train | Test | 10-fold CV | Train | Test |
| Xgboost | $0.445 \pm 0.187$ | 0.997 | 0.839 | $0.293 \pm 0.175$ | 0.997 | 0.664 |
| RF | $0.342 \pm 0.281$ | 0.908 | 0.602 | $0.256 \pm 0.298$ | 0.909 | 0.462 |
| LGBM | $0.011 \pm 0.307$ | 0.830 | 0.232 | $-0.004 \pm 0.287$ | 0.808 | 0.227 |
| SVR-rbf | $0.199 \pm 0.369$ | 0.529 | 0.294 | $0.130 \pm 0.348$ | 0.508 | 0.206 |
| SVR-linear | $0.094 \pm 0.496$ | 0.368 | 0.132 | $-0.033 \pm 0.363$ | 0.279 | 0.107 |

**Table S2.** R^2^ values for the prediction of DEmax using different ML algorithms.

|  | With mutation information |  |  | Without mutation information |  |  |
| --- | --- | --- | --- | --- | --- | --- |
|  | 10-fold CV | Train | Test | 10-fold CV | Train | Test |
| Xgboost | $0.453 \pm 0.192$ | 0.997 | 0.830 | $0.324 \pm 0.120$ | 0.997 | 0.482 |
| RF | $0.324 \pm 0.297$ | 0.907 | 0.614 | $0.218 \pm 0.321$ | 0.909 | 0.462 |
| LGBM | $0.028 \pm 0.326$ | 0.838 | 0.355 | $-0.016 \pm 0.323$ | 0.830 | 0.317 |
| SVR-rbf | $0.193 \pm 0.371$ | 0.542 | 0.302 | $0.124 \pm 0.348$ | 0.518 | 0.210 |
| SVR-linear | $-0.144 \pm 0.545$ | 0.353 | 0.160 | $-0.074 \pm 0.386$ | 0.258 | 0.176 |

**Table S3.** R^2^ values for the prediction of DE168 using different ML algorithms.

|  | With mutation information |  |  | Without mutation information |  |  |
| --- | --- | --- | --- | --- | --- | --- |
|  | 10-fold CV | Train | Test | 10-fold CV | Train | Test |
| Xgboost | $0.394 \pm 0.220$ | 0.994 | 0.741 | $0.184 \pm 0.333$ | 0.994 | 0.633 |
| RF | $0.300 \pm 0.309$ | 0.905 | 0.665 | $0.192 \pm 0.311$ | 0.901 | 0.456 |
| LGBM | $-0.005 \pm 0.333$ | 0.859 | 0.370 | $-0.042 \pm 0.374$ | 0.797 | 0.243 |
| SVR-rbf | $0.194 \pm 0.342$ | 0.525 | 0.301 | $0.149 \pm 0.315$ | 0.496 | 0.219 |
| SVR-linear | $-0.105 \pm 0.582$ | 0.382 | 0.118 | $-0.081 \pm 0.407$ | 0.247 | 0.161 |

## Notes

### Competing Interest Statement

The authors have declared no competing interest.

